# AlphaFold-Multimer reveals diverse cyclin-CDK substrate docking interactions

**DOI:** 10.64898/2026.06.29.735189

**Authors:** Sarah Willich, Nitin Kapadia, Paul Nurse

## Abstract

Cyclin-dependent kinases (CDKs) control eukaryotic cell-cycle progression by phosphorylating specific substrates with substrate recognition often involving cyclin-specific docking interactions. However, in minimal cell cycle control systems driven by a single cyclin-CDK complex, how docking interactions contribute to the differential timing of substrate phosphorylation remains unclear. Here, we used AlphaFold-Multimer to systematically predict interactions between the fission yeast mitotic cyclin-CDK fusion Cdc13-L-Cdc2 and its known *in vivo* CDK substrates. We found that many substrates are predicted to interact with the cyclin hydrophobic patch, and have identified a previously uncharacterised docking motif, [FVIPWGLAM](x)xER[LMV] (ERL motif), with features consistent with an atypical RxL motif. We show that ERL motifs can functionally substitute for canonical RxL motifs to promote phosphorylation of a model CDK substrate by Cdc13-Cdc2, while the S-phase cyclin-CDK Cig2-Cdc2 was found to preferentially phosphorylate substrates containing canonical RxL motifs. Finally, we investigated whether Cdc13-L-Cdc2 is predicted to preferentially bind DNA replication substrates over mitotic substrates but found no evidence of differential binding. These results reveal diversity in cyclin-CDK substrate recognition beyond established docking motifs.

## Introduction

Eukaryotic cells must ensure they undergo the stages of the cell cycle in the correct order, such as replicating their DNA in S-phase before segregating their chromosomes in mitosis. Progression through the cell cycle is controlled by cyclin-dependent kinases (CDKs), which initiate and trigger cell cycle events by phosphorylating their substrates. Biochemical studies in budding yeast and metazoans have led to a model in which distinct cyclin-CDK complexes possess unique specificities for different substrate subsets (Bhaduri and Pryciak, 2011; Brown et al., 1999; Kõivomägi et al., 2011; Loog and Morgan, 2005; Pagliuca et al., 2011; Peeper et al., 1993), suggesting that expressing different cyclin-CDKs at different stages of the cell cycle can form a basis for ensuring that critical CDK substrates are phosphorylated at the correct time. This model, therefore, depends on understanding how cyclin-CDK complexes interact with their substrates.

The interaction of cyclin-CDK with some of its substrates has been shown to be mediated through short linear protein motifs (SLiMs). RxL motifs (full sequence [RK]xL(x) Φ with Φ representing a hydrophobic amino acid) are the best studied SLiMs in CDK substrates. RxL motifs bind to a conserved hydrophobic patch on A-, E- and B-type cyclins (Brown et al., 1995). G1/S CDK substrates commonly contain RxL motifs, which play a role in their phosphorylation by S-cyclin-CDK (Brown et al., 1999; Kumagai et al., 2011; Örd et al., 2025; Russo et al., 1996; Wilmes et al., 2004). Additionally, in *S. cerevisiae*, the hydrophobic patch of the mitotic cyclin Clb2 specifically recognises LxF motifs, while the hydrophobic patch of the G2 cyclin Clb3 uniquely interacts with PxF motifs (Örd et al., 2019; Örd et al., 2020). Moreover, the *S. cerevisiae* S-phase cyclins Clb5 and Clb6 engage with NLxxxL motifs in addition to RxL motifs (Faustova et al., 2021). Taken together, these observations suggest that interactions with SLiMs are often cyclin-specific, thereby potentially allowing different cyclins to phosphorylate distinct subsets of substrates and fine-tune the timing of substrate phosphorylation.

However, different cyclins are not always essential for accurate eukaryotic cell cycle progression. In the fission yeast *Schizosaccharomyces pombe,* a single genetically engineered cyclin-CDK chimera between the mitotic B-type cyclin Cdc13 and the CDK Cdc2 (Cdc13-L-Cdc2) can substitute for the four cyclin-CDK complexes usually present during the mitotic cell cycle and the six acting during meiosis (Coudreuse and Nurse, 2010; Gutiérrez-Escribano and Nurse, 2015). Additionally, in budding yeast, the mitotic cyclin Clb2 alone can promote S-phase as well as mitosis (Ercan et al., 2021). In metazoans, studied using *Xenopus* egg extracts, the mitotic cyclin, Cyclin B1, can promote DNA replication and chromosome segregation (Moore et al., 2003; Prokhorova et al., 2003).

To date, SLiMs have been studied primarily in systems containing multiple cyclin-CDK complexes, where cyclin-specific docking is thought to confer substrate specificity. However, there is a lack of systematic studies examining SLiM-mediated interactions in the context of a single cyclin-CDK complex driving the entire cell cycle. The minimal *Schizosaccharomyces pombe* system, therefore, provides an opportunity to disentangle the contribution of SLiMs to substrate recognition independently of cyclin specificity.

Here, we used AlphaFold-Multimer (Evans et al., 2021) to predict interactions between over 180 CDK substrates identified in *S. pombe* (Swaffer et al., 2016) and a fusion cyclin-CDK Cdc13-L-Cdc2 (Coudreuse and Nurse, 2010). Using SLiMFinder (Davey et al., 2010), we identified a novel CDK-substrate interaction motif, [FVIPWGLAM](x)xER[LMV] (ERL motif), a potential variation of the canonical RxL motif, and validated that this motif could function in place of canonical RxL motifs in a model CDK substrate. We found that the mitotic cyclin-CDK Cdc13-Cdc2 phosphorylates a model substrate with RxL and ERL motifs with equal efficiencies, and in contrast, that the S-phase cyclin-CDK Cig2-Cdc2 preferentially phosphorylates a model substrate containing RxL motifs, indicating for the first time that different cyclins in *S. pombe* can exhibit some distinct motif preferences. We also determined that ERL motifs may be conserved in human cells to mediate CDK-substrate interactions. We further asked whether preferential binding to Cdc13-L-Cdc2 underlies the temporal order of CDK substrate phosphorylation. However, we found no association between predicted binding to Cdc13-L-Cdc2 and phosphorylation timing during the cell cycle.

## Results

### CDK substrates are predicted to bind to the Cdc13 hydrophobic patch

To systematically investigate how SLiMs contribute to substrate recognition by a single cyclin-CDK complex, we used AlphaFold-Multimer (Evans et al., 2021), specifically AlphaPulldown (Yu et al., 2023), to predict Cdc13-L-Cdc2 interactions with the CDK substrates identified in *S. pombe* (n = 182) (Swaffer et al., 2016). Predicted aligned error (PAE) is a metric that estimates AlphaFold’s confidence in the relative positions of amino acid residues (Bryant et al., 2022). A lower PAE reflects a higher confidence in the relative position of two amino acid residues. We deemed a CDK substrate to be predicted to bind to Cdc13-L-Cdc2 if it fulfilled the following criteria: 1) The substrate had residues with a PAE value of <15 with Cdc13-L-Cdc2, a PAE cut-off used by other high-throughput protein-protein interaction screens (Schmid and Walter, 2025; Sifri et al., 2023), and 2) those residues were less than 8 Å away from Cdc13-L-Cdc2 (Gromiha and Selvaraj, 1999) (Fig. 1A,B).

**Figure 1:**
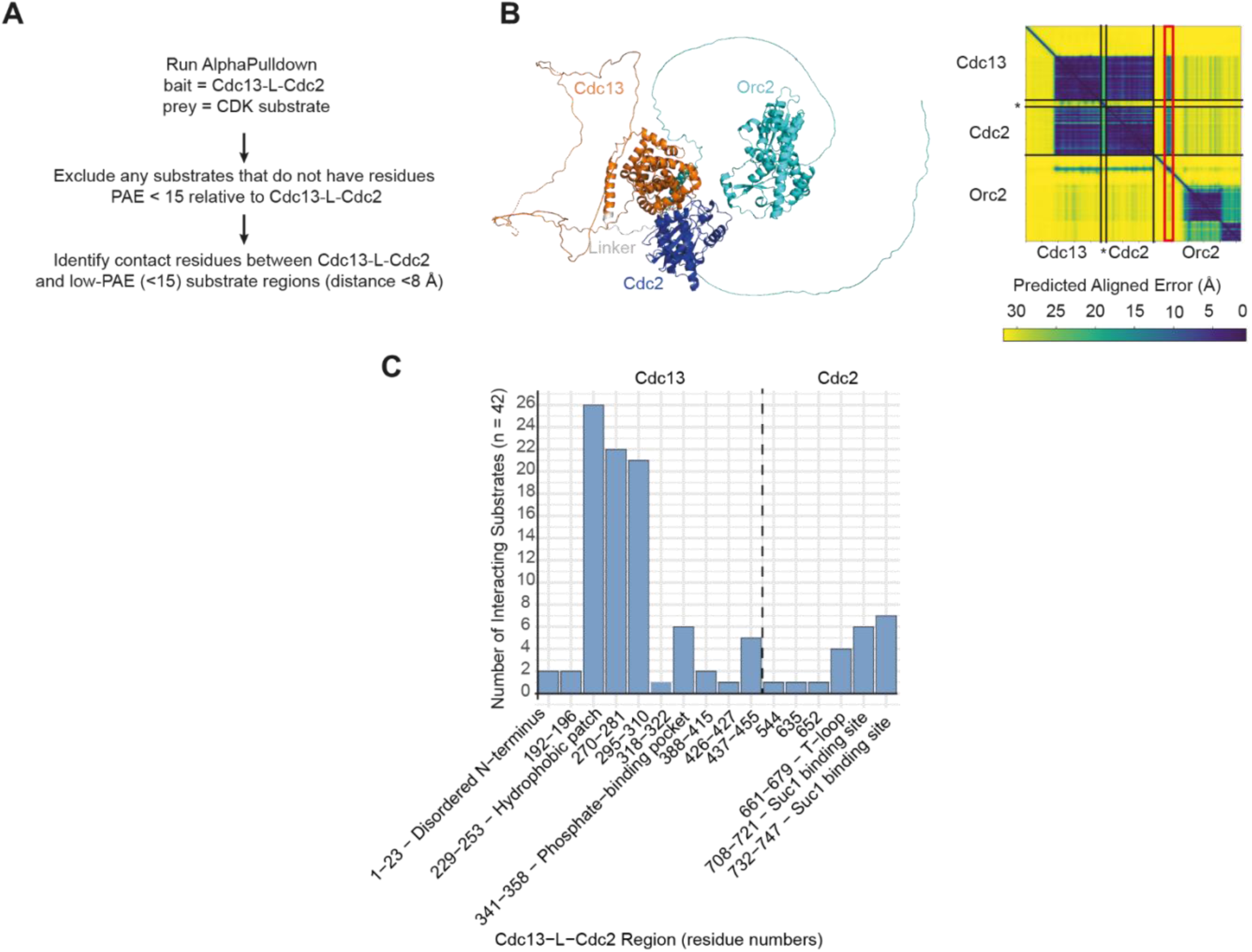
AlphaFold-Multimer Cdc13-L-Cdc2-substrate interaction screen. **A** Workflow of AlphaPulldown Cdc13-L-Cdc2-CDK substrate interaction screen. **B** Left = AlphaPulldown predicted structure of Cdc13-L-Cdc2 with Orc2. Right = Predicted Aligned Error (PAE) plot of structure shown on the left. Black lines indicate boundaries of proteins and internal fusion junctions within the chimeric Cdc13-L-Cdc2 construct. * = Linker of Cdc13-L-Cdc2. Red box highlights residues of Orc2 with a PAE < 15 relative to Cdc13-L-Cdc2. **C** Bar graph showing the number of CDK substrates predicted by AlphaPulldown to interact with regions of Cdc13-L-Cdc2. Regions are indicated by residue numbers of Cdc13-L-Cdc2. Where a function of a region of Cdc13-L-Cdc2 is known, it has been labelled as such. Black dotted line indicates the boundary between Cdc13 and Cdc2 residues in Cdc13-L-Cdc2.

Out of 182 substrates (Swaffer et al., 2016), 42 were predicted to interact with Cdc13-L-Cdc2 based on our criteria (Table S1). The other 140 substrates either did not fulfil our cut-offs for predicted interaction, or AlphaFold did not confidently resolve their potentially transient binding to Cdc13-L-Cdc2. Over half of the 42 predicted interactors (n = 26) were predicted to interact with the Cdc13 hydrophobic patch (Fig. 1C). Outside of the hydrophobic patch, six substrates were predicted to interact with the phosphate-binding pocket on Cdc13, which is conserved among B-type cyclins (Ng et al., 2025; Yu et al., 2021). The primary interaction sites predicted on Cdc2 were its T-loop (Jeffrey et al., 1995) and the interface where the CDK accessory protein Suc1 (Cks1 in other eukaryotes) also binds (Brown et al., 2015).

### An atypical RxL motif is predicted to bind to the Cdc13 hydrophobic patch

We next asked whether the substrate residues predicted to interact with the Cdc13 hydrophobic patch (and the two regions C-terminal to it, 270-281 and 295-310, Fig. 1C) were predicted to share any novel or known SLiMs. To do so in an unbiased manner, we used SLiMFinder (Davey et al., 2010), which allowed us to predict overrepresented short protein motifs in substrate residues predicted to interact with the Cdc13 hydrophobic patch.

SLiMFinder identified a shared motif, ER[LMV] (Fig. 2A), in seven substrates: Mcm10 (Fig. 2B), Bbc1, Pal1, Q09762, Tea4, Lem2, and O13695 (Fig. S1). When we generated a sequence logo of the ER[LMV] motifs and surrounding residues in the seven substrates, we found that all the substrates had at least one non-polar residue upstream of the ER[LMV] motif (Fig. 2A). Moreover, in the predicted structure of Mcm10 binding to Cdc13-L-Cdc2, the phenylalanine preceding its ERL motif plays a role in interacting with the hydrophobic patch (Fig. 2B). We concluded that this short protein motif predicted to interact with Cdc13 is [FVIPWGLAM](x)xER[LMV] (hereafter referred to as the ERL motif). We noted that in all seven predicted ERL motifs, the glutamic acid residue was not predicted to face the Cdc13 hydrophobic patch (Fig. S1). Therefore, we hypothesise that this could be an atypical RxL motif, defined by deviations from the typical RxL consensus, which have recently been systematically described by Örd et al (2025) in a high-throughput screen of human cyclin-binding peptides.

**Figure 2:**
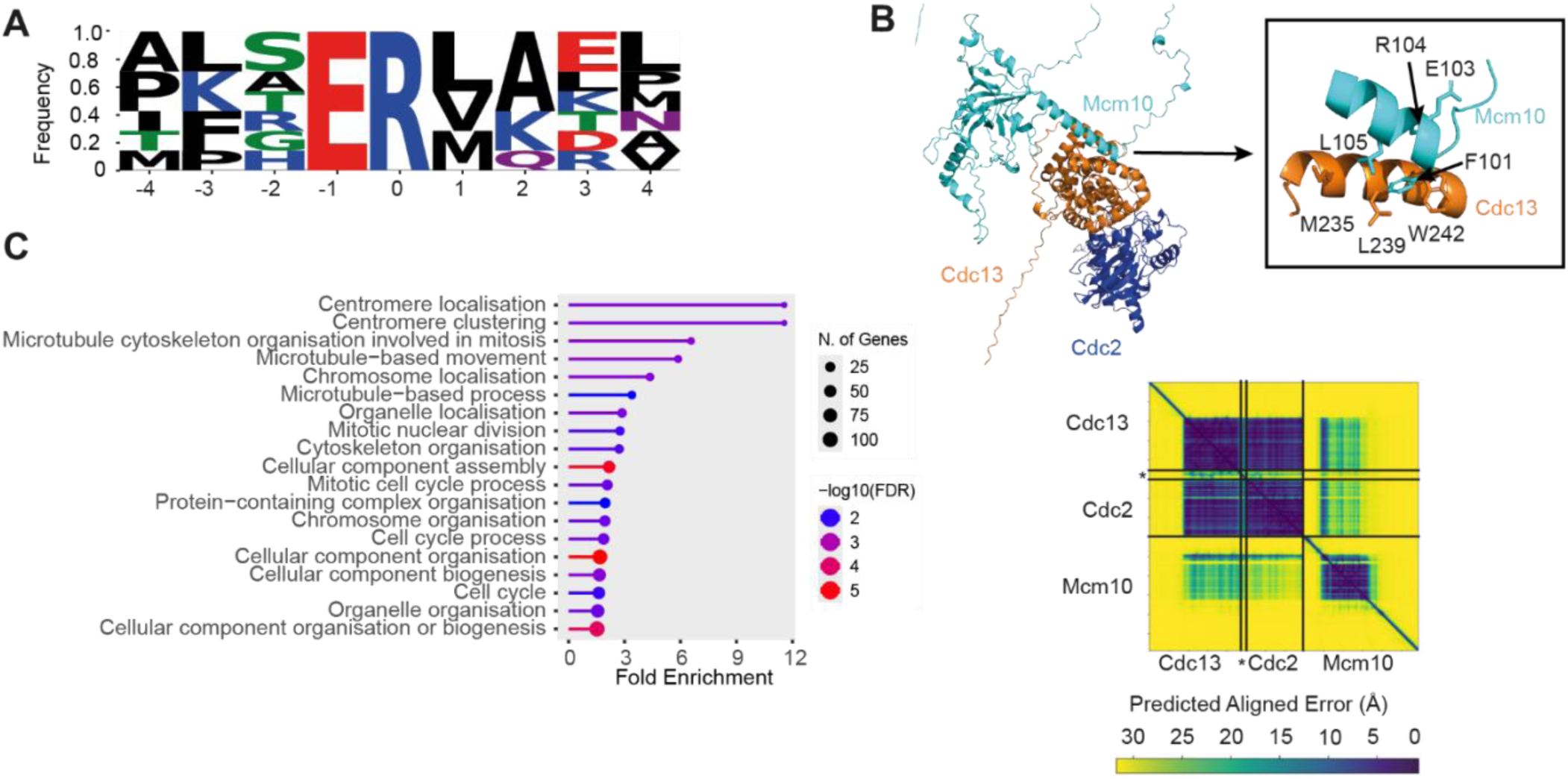
Identifying a novel short protein motif predicted to interact with the Cdc13 hydrophobic patch. **A** Filled sequence logos of ER[LMV] motifs in Mcm10, Bbc1, Pal1, Q09762, Tea4, Lem2 and O13695 predicted to bind the Cdc13 hydrophobic patch + 3 residues up- and downstream. Amino acids are coloured by physicochemical properties = positive (blue), negative (red), hydrophobic (black), hydrophilic (purple), and small, uncharged residues (green). **B** Top = Predicted structure of Cdc13-L-Cdc2 with Mcm10. Box on the right = Cdc13 hydrophobic patch (M235, L239, W242) predicted to interact with FxERL motif on Mcm10 (F101, E103, R104, L105). Bottom = corresponding PAE plot. Black lines indicate boundaries of proteins and internal fusion junctions within the chimeric Cdc13-L-Cdc2 construct.* = Linker of Cdc13-L-Cdc2. **C** GO biological process of proteins with ERL motifs in the *S. pombe* proteome. FDR = False Discovery rate.

Alongside PAE, AlphaFold uses a second confidence metric, the Predicted Local Distance Difference Test (pLDDT) (Jumper et al., 2021). The predicted structure of Mcm10 in complex with Cdc13-L-Cdc2 showed that its FxERL motif is predicted to be part of an alpha helix, which was predicted with high confidence (pLDDT 70-90) (Fig. 2B, Fig. S2A). We compared the predicted structures of the monomers for each of the seven substrates to their structures when bound to Cdc13-L-Cdc2. In the case of Mcm10 and Bbc1, the ERL motif is predicted to be part of an alpha helix in both the monomer and in the predicted interaction with Cdc13-L-Cdc2 (Fig. S2A,B). In the case of Lem2, its ERL motif was predicted to be disordered in the monomer but was alpha-helical in its predicted interaction with Cdc13-L-Cdc2, although with a low corresponding pLDDT (Fig. S2C). The low pLDDT for the Lem2 alpha helix could reflect a coupled folding and binding mechanism, in which the motif is natively disordered in isolation but adopts a helical conformation upon interaction with the Cdc13-L-Cdc2 (Alderson et al., 2023). This suggests that ERL motifs are predicted to adopt alpha-helical conformations when bound to the Cdc13 hydrophobic patch.

Surprisingly, SLiMFinder did not identify previously characterised cyclin hydrophobic patch-interacting motifs such as RxL, NLxxL, LxF or PxF (Faustova et al., 2021; Örd et al., 2019; Örd et al., 2020; Russo et al., 1996) among the substrate residues predicted to bind to Cdc13. However, we noted that the DNA replication initiation protein Sld3 was predicted to have a conserved RxL motif near the Cdc13 hydrophobic patch (Fig. S3), which has been shown to be important for the phosphorylation of its human analogue Treslin (Kumagai et al., 2011).

### ERL motifs are found in putative CDK substrates

To investigate whether ERL motifs are found in proteins beyond known CDK substrates, we searched the entire *S. pombe* proteome to identify proteins containing ERL motifs ([FVIPWGLAM](x)xER[LMV]) in accessible parts of proteins using SLiMSearch 4 (Krystkowiak and Davey, 2017). We identified 214 ERL motifs across 202 proteins out of 5,147 proteins in the *S. pombe* proteome (Table S2). To investigate the biological roles of these proteins, we performed GO enrichment (Fig. 2C). This analysis showed that proteins with ERL motifs are enriched in processes including centromere localisation, microtubule-based processes, and the mitotic cell cycle. Of these 202 proteins, 17 were identified as CDK substrates by Swaffer et al. (2016), and 14 more have recently been shown to be phosphorylated by CDK (Curran et al., 2025).

We also investigated whether some of these proteins could be unidentified CDK substrates by assessing whether they have putative CDK phosphosites. CDK phosphosites are commonly found in disordered regions of proteins (Holt et al., 2009). Of the 202 ERL motif-containing proteins, 117 (57.9 %) have putative CDK phosphosites in intrinsically disordered regions of the protein (S/TP in IUPred3 cut-off > 0.5) compared to 33.8 % of the total proteome (1,737/5,147) (chi-squared test p = 1.4e-12), indicating that ERL-motif-containing proteins are enriched for putative disordered CDK phosphosites relative to the proteome background. These findings suggest that ERL motifs are associated with proteins involved in the mitotic cell cycle and are putative CDK substrates.

### ERL motifs can substitute for RxL motifs in a model CDK substrate

We next investigated whether ERL motifs function as CDK-substrate interaction motifs *in vivo*. We used the single-cell CDK sensor NucCDK, which translocates from the nucleus to the cytoplasm upon phosphorylation by CDK, allowing sensor phosphorylation to be quantified by its cytoplasmic-to-nuclear ratio in time-lapse microscopy (Kapadia and Nurse, 2025) (Fig. 3A). NucCDK sensor-readout increases gradually over the course of G2 before a rapid increase corresponding to mitotic CDK activity (Kapadia and Nurse, 2025) (Fig. 3B, Fig. S4A).

**Figure 3:**
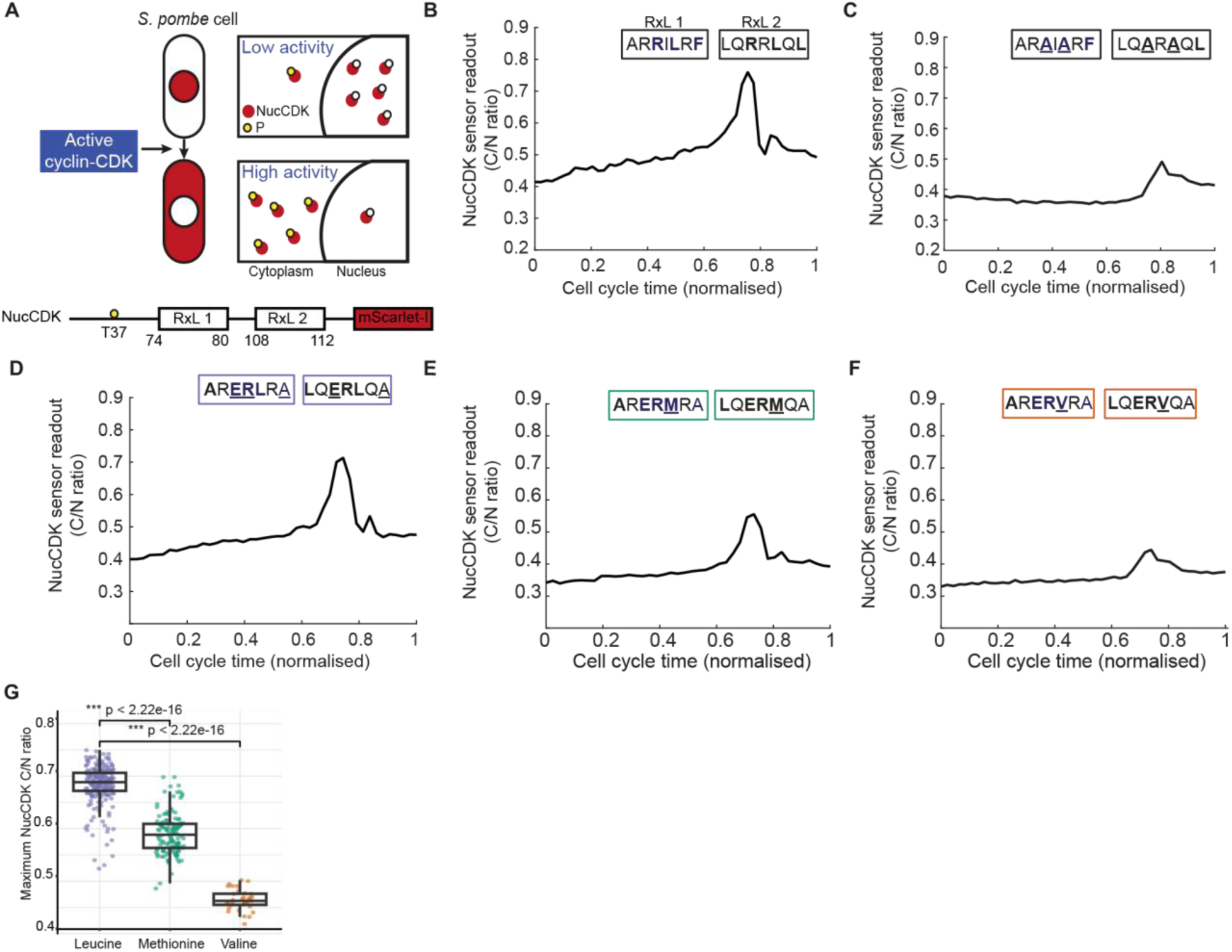
ERL motifs can replace RxL motifs in the NucCDK sensor. **A** Top = Schematic showing the principle of NucCDK sensor. Bottom = schematic representation of NucCDK sensor. T37 = essential phosphorylation site for export of the sensor. RxL 1 and RxL 2 represent the two RxL motifs found in NucCDK. **B-F** Representative profile of NucCDK-mScarlet-I showing the cytoplasmic-to-nuclear ratio of mean intensity (C/N). Above = amino acid sequence of **B** RxL motifs and **C-F** their mutations. Bold characters indicate motif. Underlined characters highlight mutations relative to previous motif. 1 of 2 biological replicates. **B** NucCDK with intact RxL motifs. **C** RxL motifs mutated to AxA. **D** RxL motifs mutated to ERL motif. **E** RxL motifs mutated to ERM motif. **F** RxL motifs mutated to ERV motif. **G** Boxplot showing the median maximum NucCDK C/N ratio in cells shown in D-F. Only cells that showed at least a change of 0.1 in their C/N ratio were included in this analysis. Leucine n = 223, Methionine n = 130, Valine n = 33. Results (p-value) of a two-sided Wilcoxon rank-sum test (Mann-Whitney U test) are displayed on the plot.

NucCDK has two RxL motifs (RxLxF and RxLxL), (Fig. 3A), and we found that they are required for its phosphorylation, since mutating both RxL motifs (AxAxF and AxAxL) abolished most of the phosphorylation of NucCDK as measured by the change in its C/N ratio (Fig. 3C, Fig. S4B). This suggests that NucCDK relies on its interaction with Cdc13 via one or both of its RxL motifs for phosphorylation.

We investigated whether substituting the RxL motifs for ERL motifs would preserve the NucCDK sensor phosphorylation pattern or abolish it. Replacing RxL with ERL motifs (AxERLxA and LxERLxA) resulted in a similar phosphorylation pattern in the sensor (Fig. 3D, Fig. S4C). On the other hand, while ERM and ERV showed a similar overall pattern of phosphorylation, the amplitude of their phosphorylation was significantly lower than that of the leucine variant of the motif (Fig. 3E-G). The ERV motif version appears identical to the scenario in which the RxL motifs are completely mutated and abolished (Fig. 3C&F, Fig. S4B&E). Therefore, we concluded that leucine provides a stronger interaction than methionine and that valine barely allows the phosphorylation of the sensor. These results suggest that the ERL motif can be classified as an atypical RxL motif, a category previously described by Örd et al. (2025).

We found that the interaction of the NucCDK RxL or ERL motif versions with Cdc13-Cdc2 predicted using AlphaFold-Multimer (Evans et al., 2021) indicated that one of the RxL motifs and one of the ERL motifs were both predicted to interact with Cdc13 as an alpha helix (Fig. S5). Collectively, these results demonstrate that ERL motifs can functionally replace RxL motifs in a CDK substrate, with the terminal leucine residue being critical for efficient docking and subsequent phosphorylation.

### S-phase cyclin-CDK preferentially phosphorylates model substrate with RxL over ERL motifs

In the GO analysis of ERL motif instances in the *S. pombe* proteome, proteins involved in mitosis rather than DNA replication were enriched. Since many cyclin-binding SLiMs display cyclin specificity (Faustova et al., 2021; Kõivomägi et al., 2011; Örd et al., 2020), we next asked whether the S-phase cyclin Cig2 could promote the phosphorylation of an ERL-containing substrate or if the ERL motif is uniquely recognised by Cdc13.

Because Cig2 is predominantly present during S-phase in contrast to Cdc13 (Mondesert et al., 1996), we used a NucCDK variant containing a serine phosphoacceptor (SPAK rather than TPAK), which is phosphorylated at the G1/S transition (Liku et al., 2005), as a model substrate (Kapadia and Nurse, 2025). We then replaced its canonical RxL docking motif with ERL motifs and compared the nuclear-to-cytoplasmic translocation of both the RxL- and ERL-motif sensors.

To eliminate the influence of cyclins other than Cig2 on the phosphorylation of the sensor, we conducted the experiment in strains in which the other G1/S cyclins (Puc1 and Cig1) were absent (*puc1Δ cig1Δ*), and Cdc13 was under the expression of the thiamine-repressible promoter nmt41. First, we synchronised cells in G1 by nitrogen starvation, and one hour before releasing them from the arrest, we split the cells and either omitted or added thiamine to leave on or shut off *cdc13* expression. We then monitored the translocation of the sensor in single cells under each condition using time-lapse microscopy (Fig. 4A).

**Figure 4:**
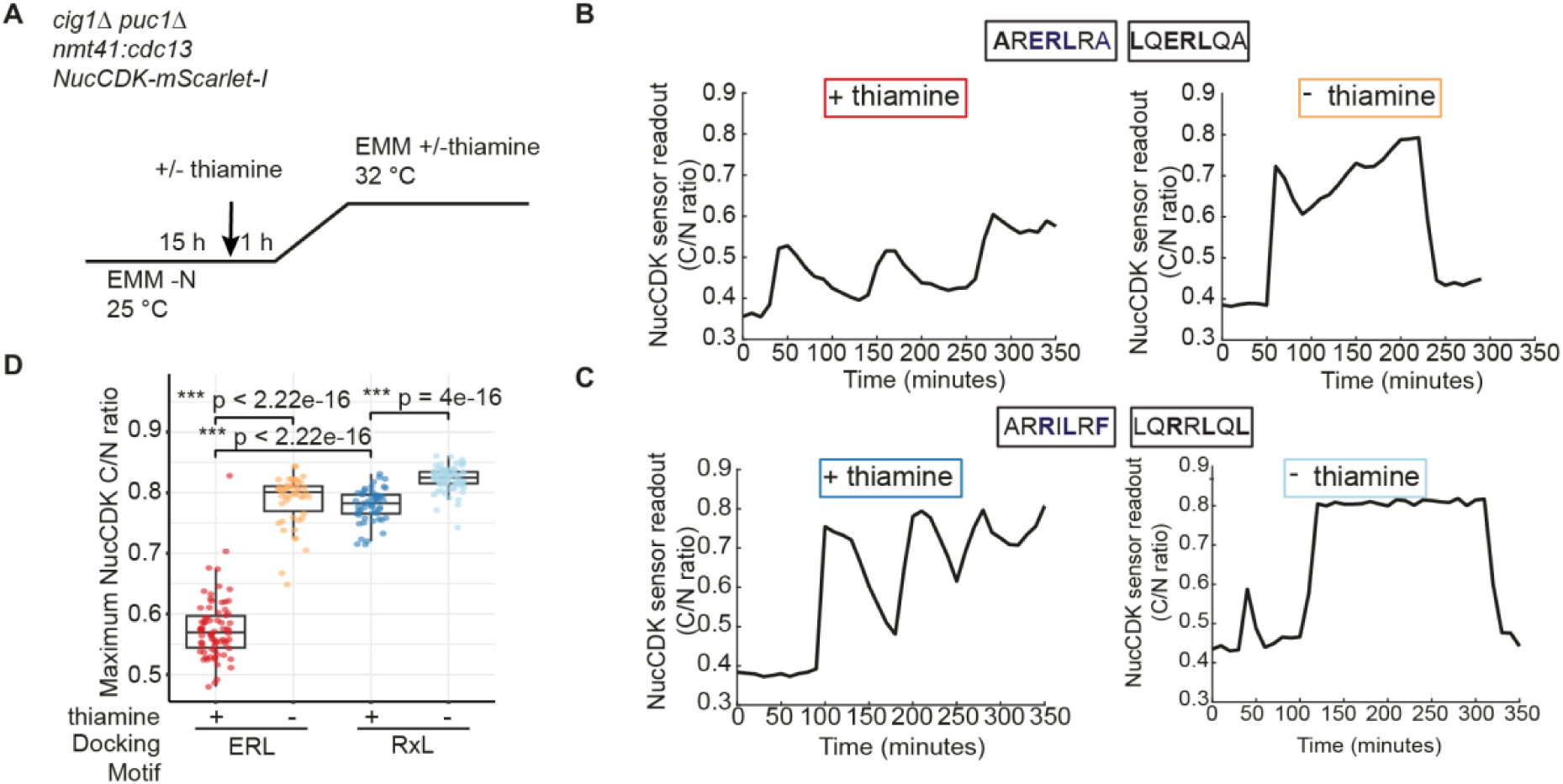
S-cyclin-CDK preferentially phosphorylates RxL over ERL motifs. **A** Schematic representation of the time-lapse microscopy experiment. Genotype of strains used is displayed. NucCDK-mScarlet-I denotes either ERL or RxL motif version of the sensor. **B-C** Representative profile of NucCDK-mScarlet-I ERL (**B**) or RxL (**C**) motif version showing the cytoplasmic to nuclear ratio of mean intensity (C/N) of cells with (left) and without (right) thiamine over time after release from G1 arrest. **D** Boxplot showing the median maximum NucCDK C/N ratio in cells shown in **B-C**. ERL motif + thiamine n = 74, ERL motif - thiamine n = 50, RxL motif + thiamine n = 53, RxL motif - thiamine n = 78. Results (p-value) of a two-sided Wilcoxon rank-sum test (Mann-Whitney U test) are displayed on the plot.

Notably, in both sensor versions, when Cdc13 expression was repressed (+ thiamine), the sensor displayed a periodic pattern of phosphorylation (Fig. 4B&C, Fig. S6, + thiamine). This is consistent with the observation that cells lacking Cdc13 undergo endoreduplication cycles dependent on Cig2 (Ayté et al., 2001; Fisher & Nurse, 1996).

While the ERL motif version of NucCDK did get phosphorylated by Cig2-Cdc2 in the absence of *cdc13* expression (+ thiamine), the maximum C/N ratio displayed by the sensor was lower than in the presence of Cdc13 (- thiamine) (Figure 4B-D, Fig. S6). This was specific to the ERL motif version of NucCDK and was not due to lower Cig2-Cdc2 activity compared to Cdc13-Cdc2, since the RxL motif version of NucCDK showed a similar maximum C/N ratio in the presence or absence of Cdc13 (Figure 4C&D). These results suggest that Cig2-Cdc2 preferentially phosphorylates the RxL motif version of NucCDK over the ERL motif version.

### ERL motifs are found in human proteins

Since we established a possible role for ERL motifs in regulating CDK-substrate interactions in *S. pombe*, we aimed to determine whether this was potentially conserved. First, we sought to understand which proteins in the human proteome have putative ERL motifs and what their functions are. Out of 20,659 proteins in the human proteome, we identified 1,749 ERL motif occurrences in 1,492 proteins using SLiMSearch 4 (Krystkowiak and Davey, 2017) (Table S3), 1,144 of which (76.6 %) have putative CDK phosphosites (S/TP) in intrinsically disordered regions of the protein (IUPred3 cut-off > 0.5), compared to 50.9 % (10,514/20,659) of the total proteome (chi-squared test p = 1.1e-82). Moreover, 98 (out of 548) of these proteins were previously identified as CDK1 and 10 (out of 180) as CDK2 substrates in human cells (Chi et al., 2008; Petrone et al., 2016). Additionally, in fixed, permeabilised human cells, 322 of the ERL-motif-containing substrates were phosphorylated by Cyclin B1-CDK1 (Al-Rawi et al., 2023).

Moreover, as seen in *S. pombe*, ERL motif-containing proteins were enriched for biological processes involving microtubule-based processes and cytoskeleton organisation (Fig. 5A). CDKs are known to phosphorylate proteins at the centrosome and microtubule components involved in spindle assembly in human cells (Blangy et al., 1995; Nigg and Raff, 2009; Ookata et al., 1997).

**Figure 5:**
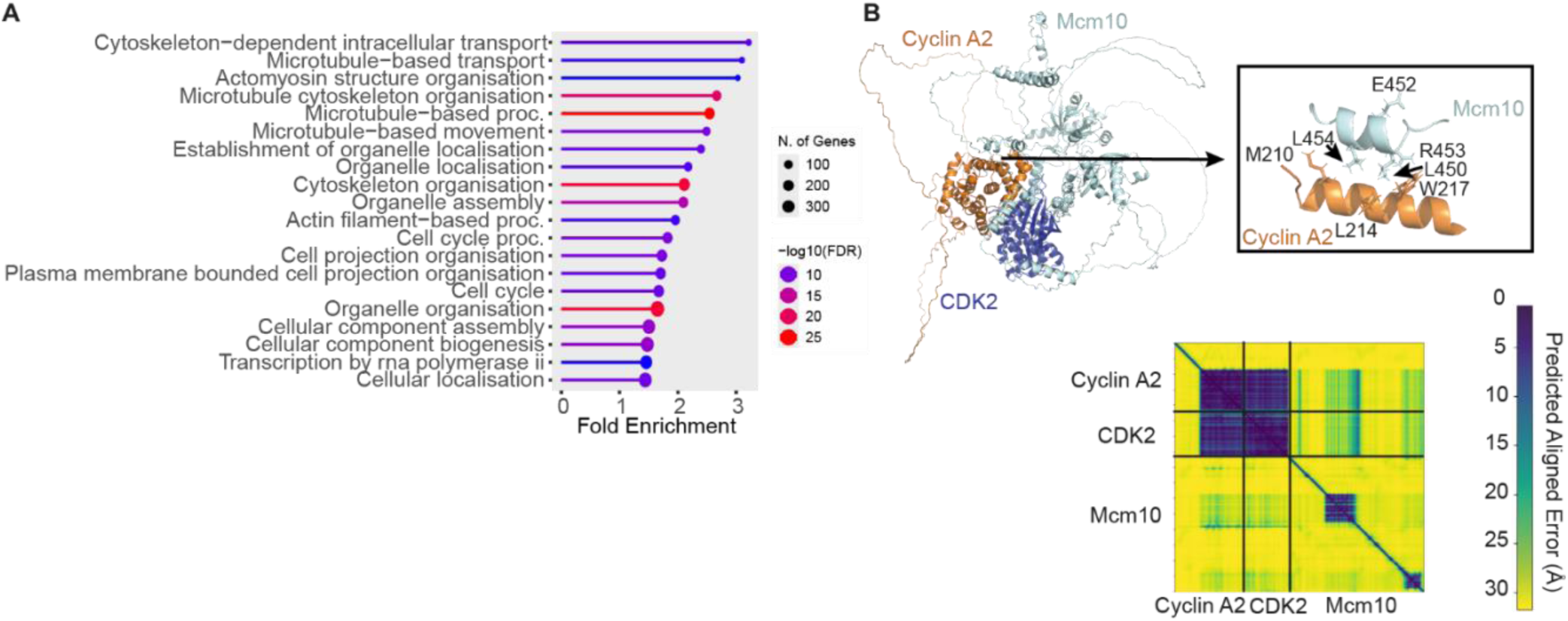
ERL motifs are found in human proteins. **A** GO biological process of proteins with ERL motifs in the human proteome. FDR = False Discovery rate. **B** Predicted AlphaFold-Multimer structure of human Cyclin A2-CDK2 with MCM10. Box on the right = Cyclin A2 hydrophobic patch (M210, L214, W217) predicted to interact with LxERL motif on MCM10 (L450, E452, R453, L454). Bottom = corresponding PAE plot. Black lines indicate boundaries of proteins.

Finally, we investigated whether the predicted interaction between the conserved DNA replication protein Mcm10 and cyclin is evolutionarily conserved in humans. Using AlphaFold-Multimer (Evans et al., 2021), we predicted the interaction between human MCM10 and Cyclin A2-CDK2. We chose Cyclin A2 instead of Cyclin E because Cyclin E functions predominantly at the G1/S transition and can be partially compensated for by other cyclin-CDK complexes (Geng et al., 2003), whereas Cyclin A2 is the major cyclin driving S-phase progression (Pagano et al., 1992). As seen in *S. pombe* Mcm10, human MCM10 was found to engage the Cyclin A2 hydrophobic patch through an alpha-helical ERL motif (LxERL specifically) (Fig. 5B). Ultimately, these results establish ERL motifs as a potentially conserved feature of CDK-substrate recognition from yeast to humans.

### CDK-substrate interactions and the order of the cell cycle

A key unresolved question is how a single cyclin-CDK complex can drive a temporally ordered cell cycle. In *S. pombe* and fixed permeabilised human cells, DNA replication substrates (early substrates) are phosphorylated when CDK activity is low, whereas G2 and mitotic substrates (mid and late substrates, respectively) require higher CDK activity to be phosphorylated (Al-Rawi et al., 2023; Swaffer et al., 2016). DNA replication (early) substrates have been proposed to more readily interact with cyclin-CDK than mitotic (late) substrates (Al-Rawi et al., 2023; Swaffer et al., 2016), to explain their phosphorylation at low CDK activity.

To investigate whether this differential affinity is reflected in our AlphaFold predictions, we examined the proportions of early, mid, and late substrates, as well as substrates with unknown cell cycle dynamics (Swaffer et al., 2016), predicted to bind the Cdc13 hydrophobic patch (Table S1). Three out of 16 early substrates (18.75 %), three out of 13 mid substrates (23 %), 15 out of 165 late substrates (9 %) and seven out of 60 substrates with unknown cell cycle dynamics (11.6 %) were predicted to bind the hydrophobic patch (Fig. 6). Therefore, the proportions of predicted hydrophobic patch binders did not differ significantly between early, mid, and late substrates. However, AlphaFold does not provide information on the binding efficiency or affinity of any predicted protein-protein interaction, so, it is possible that early substrates interact more strongly with the hydrophobic patch than late substrates do. However, the hydrophobic patch of both Cdc13 and Cig2 is dispensable for S-phase progression in *S. pombe* (Basu *et al*, 2020; Roberts *et al*, 2024). Therefore, it seems unlikely that this interaction site on Cdc13-L-Cdc2 discriminates between substrates to enforce the specific temporal order of their phosphorylation during the cell cycle.

**Figure 6:**
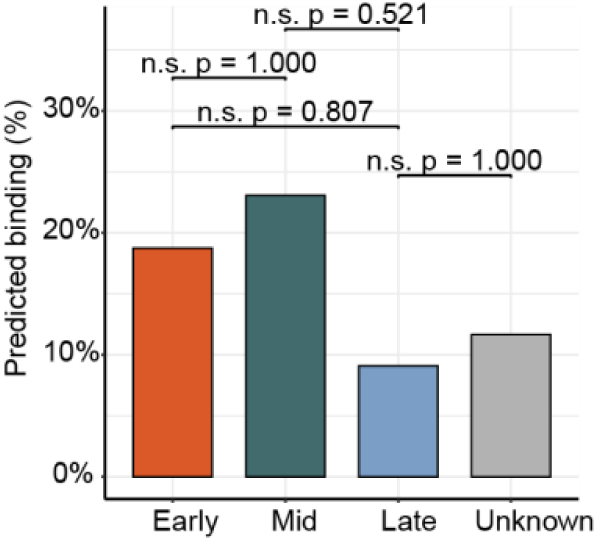
Substrate phosphorylation timing and predicted interaction with the Cdc13 hydrophobic patch. Proportion of early, mid and late substrates, and substrates with unknown cell cycle dynamics predicted to bind to the Cdc13 hydrophobic patch. Results (p-values) of pairwise Fisher’s exact tests are shown. n. s. = not significant. Cell cycle dynamics as described in Swaffer et al. (2016). If a substrate had multiple phosphosites with different cell cycle dynamics, we assigned the most common cell cycle timing to it (for a site with an unknown phosphorylation time, having an early, mid or late site took precedence).

## Discussion

In this work, we performed an AlphaFold-Multimer screen to systematically investigate how CDK substrates (Swaffer et al., 2016) are predicted to interact with a single cyclin-CDK complex (Cdc13-L-Cdc2) in the fission yeast, *S. pombe*. A notable observation from our screen is that only 42 of 182 known CDK substrates (Swaffer et al., 2016) were predicted to interact with Cdc13-L-Cdc2. Many CDK substrates may be phosphorylated efficiently through proximity and intrinsic sequence features alone, without requiring stable docking interactions. The substrates we do detect may therefore represent those with the strongest or conserved docking contacts, rather than a complete picture of the interactome.

We found that AlphaFold predicted that the majority of substrates interact with the Cdc13 hydrophobic patch. To our surprise, in our screen, we did not identify an overrepresentation of previously identified SLiMs shown to interact with cyclin-CDK (Faustova et al., 2021; Örd et al., 2019; Örd et al., 2020; Russo et al., 1996) among the substrate residues predicted to bind the hydrophobic patch. However, a recent large-scale investigation into human cyclin-peptide binding has revealed a more complex landscape, characterised by significant sequence diversity among peptides that interact with human cyclins, suggesting that the ‘rules’ for cyclin docking may be less rigid than previously thought (Örd *et al*., 2025).

We also found that six substrates were predicted to interact with the residues of Cdc2, where Cks1 is known to bind. This result was initially surprising because the Cks1-binding interface has been considered to be a site exclusively for regulatory protein recruitment rather than a direct docking site for substrates. Cdc25 was one of the substrates predicted to bind to this interface (Table S1). A recent Cryo-EM structure has shown that human Cdc25A interacts with CDK2 at the Cks1-binding interface (Rowland et al., 2024), showing that this interface is not exclusive to Cks1. Moreover, in the absence of Suc1, Cdc25 shows a 3-fold increase in its association with Cdc2 in *S. pombe* (Curran et al., 2025).

We identified a novel Cdc13 interaction motif, the ERL motif. The upstream non-polar residues that are part of the ERL motif were also commonly found in human cyclin-binding peptides (Örd et al., 2025). However, variations of the basic residue (R/K) found in RxL motifs (E in ERL motifs) were rarely observed in human cyclin-binding peptides (Örd et al., 2025). We found that ERL motifs were predicted in alpha-helical regions of substrates or to become alpha-helical upon binding to the Cdc13 hydrophobic patch. The only other helical motif described to bind to a B-type cyclin is found on MAD1 (Jackman et al., 2020). Our findings suggest that helical binding might be more prevalent than previously thought. Rather than being restricted to intrinsically disordered regions, docking interactions may also occur within structured, alpha-helical elements of substrates.

We found ERL motifs to be enriched in proteins involved in microtubule-based processes in *S. pombe* and humans. We speculate that docking interactions may enhance substrate phosphorylation efficiency in the cytoplasm since cyclin-CDK is less concentrated there than in the nucleus in *S. pombe* (Alfa et al., 1989; Curran et al., 2022). This hypothesis is consistent with the observation that a Cdc13 hydrophobic patch mutant cannot initiate mitosis due to insufficient phosphorylation of cytoplasmic and centrosomal substrates (Basu et al., 2020). The potential importance of docking interactions in overcoming spatial constraints on cyclin-CDK activity may extend beyond fission yeast, as cyclin-CDK complexes are also subject to dynamic subcellular compartmentalisation in human cells. Human Cyclin A2 is largely nuclear, and Cyclin B1-CDK1 accumulates in the cytoplasm during G2 (Pines and Hunter, 1991) but remains inactive until mitotic entry, when it undergoes a switch-like activation associated with rapid nuclear import (Gavet and Pines, 2010; Maryu and Yang, 2022).

We found that the S-phase cyclin Cig2 shows a preference for RxL over ERL motifs, which was unexpected since Cig2 and Cdc13 can phosphorylate many of the same substrates (Basu et al., 2022). However, some substrates show preferential phosphorylation by Cdc13 (Basu et al., 2022), consistent with the possibility that some docking differences can contribute to cyclin-specific substrate selection to fine-tune substrate phosphorylation, even though overall substrate overlap is substantial. These results are consistent with what has been found for human cyclins, with G1/S cyclins being broadly permissive for canonical RxL motif binding, whereas Cyclin B1 commonly engages atypical RxL motifs (Örd et al., 2025).

Finally, we sought to investigate how temporally ordered CDK substrate phosphorylation is achieved in cell cycles driven by a single cyclin-CDK complex. Although DNA replication substrates have been proposed to interact more efficiently with CDK than mitotic substrates (Al-Rawi et al., 2023; Swaffer et al., 2016), our analysis does not support a model in which differential engagement of the Cdc13 hydrophobic patch explains this ordering. Instead, we suggest that temporal phosphorylation is governed by differences in substrate affinity together with additional layers of regulation, such as subcellular compartmentalisation.

In summary, our work shows that CDK-substrate interactions in a minimal cyclin-CDK system are more diverse than previously appreciated. The identification of the ERL motif and the flexibility in motif usage suggest that substrate recognition involves a broader range of sequence and structural features than the canonical SLiM framework implies. We propose that docking motifs provide a layer of fine-tuning within a broader regulatory landscape - one in which subcellular localisation and additional post-translational regulatory mechanisms contribute to ordering cell cycle events. Understanding how these layers are integrated will be key to building a complete mechanistic picture of CDK-cell cycle control.

## Methods

### AlphaFold analysis

To predict Cdc13-L-Cdc2 (Coudreuse and Nurse, 2010) in complex with all substrates identified by Swaffer et al. (2016), AlphaPulldown (v0.22.3) (Yu et al., 2023) was performed (mode=pulldown, max_template_date=2050-01-01, num_cycle=3, num_predictions_per_model=1). Structure predictions outside of the screen were conducted using AlphaFold2.3 (Evans et al., 2021; Jumper et al., 2021). Predicted aligned error (PAE) and predicted LDDT (pLDDT) data were extracted from JSON files using AlphaPickle (Arnold, 2021). High-throughput structure analysis was performed using custom Python3 scripts. Substrate residues of interest were defined to have a PAE of less than 15 against at least ten residues of Cdc13-L-Cdc2 from the highest confidence prediction (ranked_0). Substrate residues of interest were subsequently classed as being in contact with Cdc13-L-Cdc2 residues if they were predicted to be 8 Å or closer in the predicted structure based on the distance between the Cα atom of each residue. To identify enriched SLiMs, the SLiMFinder bioinformatics tool (Davey et al., 2010) was used to analyse substrate residues predicted to bind to a region of Cdc13-L-Cdc2 (+10 residues up- and downstream).

Protein structures presented in this work were generated in PyMOL (v3.1.6.1). Monomer structures were obtained from the AlphaFold Protein Structure Database (https://alphafold.ebi.ac.uk/) (Fleming et al., 2025). Colouring of structures corresponding to pLDDT values was performed using alphafold_coloring.py (https://github.com/ailienamaggiolo/alphafold_coloring) (Fig. S2 & S5B,D).

### Proteome-wide motif screen and GO enrichment

SLiMSearch 4 (Krystkowiak and Davey, 2017) was used to identify ERL motif-containing proteins in the *S. pombe* and human proteome (default settings). GO enrichment analysis was performed using ShinyGo 0.85.1 (Ge et al., 2020).

### *Schizosaccharomyces pombe* genetics and cell culture

Culture conditions and media preparation for *S. pombe* were performed following standard protocols (Moreno et al., 1991). The full list of strains used in this study is provided in Table S4. Cells were cultured on solid YE4S agar (yeast extract supplemented with adenine, leucine, histidine, and uridine at 0.25 g/L each) or solid EMM4S agar (Edinburgh minimal media supplemented with adenine, leucine, histidine, and uridine at 0.25 g/L each). Liquid cultures in YE4S or EMM4S were maintained in exponential growth at temperatures of either 25 °C or 32 °C.

To induce nitrogen starvation, cells were initially grown in standard EMM. Cells were then harvested by centrifugation at 3000 g for 2 min, washed four times with nitrogen-free EMM medium, and resuspended to a final cell density of 2 x 10⁶ cells/mL in nitrogen-free EMM. The cells were subsequently incubated with shaking at 25 °C for 16 h to achieve cell cycle arrest. For adenine auxotrophic strains, adenine was added at 0.05 g/L to the arrest medium and at 0.25 g/L to the growth medium. When required for promoter repression, thiamine hydrochloride (Sigma), dissolved in water, was introduced to a final concentration of 30 μM one hour prior to releasing cells from arrest.

### Time-lapse microscopy

Live-cell imaging was performed on agarose pads. Gene Frames (Thermo Fisher Scientific) were placed side-by-side on a glass slide to create wells. Agarose (1 %) was prepared by heating 10 mg of ultrapure agarose (Invitrogen) in 1 mL of growth medium at 70 °C with mixing at 1000 rpm. 100 μL of molten agarose was added to each frame, flattened with a coverslip, and allowed to set. Once solidified, coverslips and backing were removed to expose the adhesive borders. Exponentially growing cells were pelleted at 2000 rpm for 30 s, and ∼950 μL of the supernatant was discarded. A small volume (1.5 μL) of the concentrated cells was spotted onto the agarose surface in a 3 x 4 grid. After spotting, a fresh coverslip was applied and sealed with fast-drying clear nail varnish (Boots).

Imaging was performed using a Nikon Ti2-E inverted widefield system, equipped with a 100x Plan Apochromat oil objective (NA 1.45) and a Prime sCMOS camera (Photometrics) with an Okolab environmental chamber used to maintain stage temperature at either 25 °C or 32 °C. Fluorescent signals were acquired using a SpectraX LED light engine (Lumencor). mScarlet-I was excited at 575/25 nm, with emission captured using a 641/75 nm filter (Semrock). Fields of view were selected and acquisition started within 30 min of mounting. Z-stacks were typically captured in five optical sections, spanning ±1 μm around the focal plane in 0.5 μm increments.

Imaging was automated via the Micro-Manager software (v2.0) (Edelstein et al., 2010). Timepoints were typically acquired every 5 min unless stated otherwise. Brightfield images were collected 1 μm below the default z-position (−2 μm total offset) to improve segmentation. A Perfect-Focus System (PFS) was used to maintain focal stability throughout acquisition.

### Image processing

Image processing was done using Fiji (ImageJ v2.14 or earlier) (Schindelin et al., 2012) and custom MATLAB scripts. Cell boundaries were identified from brightfield images at −1 μm z-slice and segmented using MATLAB routines (Nitin Kapadia; https://github.com/nkapadia27/Spatiotemporal-Orchestration-of-Mitosis).

Segmentation masks were manually reviewed and corrected when necessary. Fluorescence images were collapsed into a single plane via maximum projection across all five z-slices. To quantify signal intensity, the mean pixel value within each cell mask was calculated from the projected images. Cytoplasmic-to-nuclear fluorescence ratios were estimated by using the average of the brightest 15 % of pixels to divide by the remaining 85 %, based on the nuclear-to-cytoplasmic area ratio typical of *S. pombe* cells. Cell tracking was conducted using the Lineage Mapper plugin in Fiji (Chalfoun et al., 2016) in combination with MATLAB scripts (Nitin Kapadia; https://github.com/nkapadia27/Spatiotemporal-Orchestration-of-Mitosis).

## Data and resource availability

Previously described custom scripts used to analyse fluorescence time-lapse data can be found at https://github.com/nkapadia27/Spatiotemporal-Orchestration-of-Mitosis.

## Acknowledgements

We thank James Campbell from the Bioinformatics STP at the Francis Crick Institute for his help in running AlphaPulldown. We thank Stephane Mouilleron from the Structural Biology STP at the Francis Crick Institute for input on the AlphaFold-Multimer analysis. We thank Jessica Greenwood, Joseph Curran, Thomas Hammond and Souradeep Basu for their comments on the manuscript.

## Author contributions

S.W. and P.N. initiated the study. N.K. constructed the AxA mutant of the NucCDK sensor and characterised its phenotype. S.W. designed and performed all other experiments. S.W. performed all data analysis. S.W. and P.N. wrote the manuscript.

---

**Figure S1:**
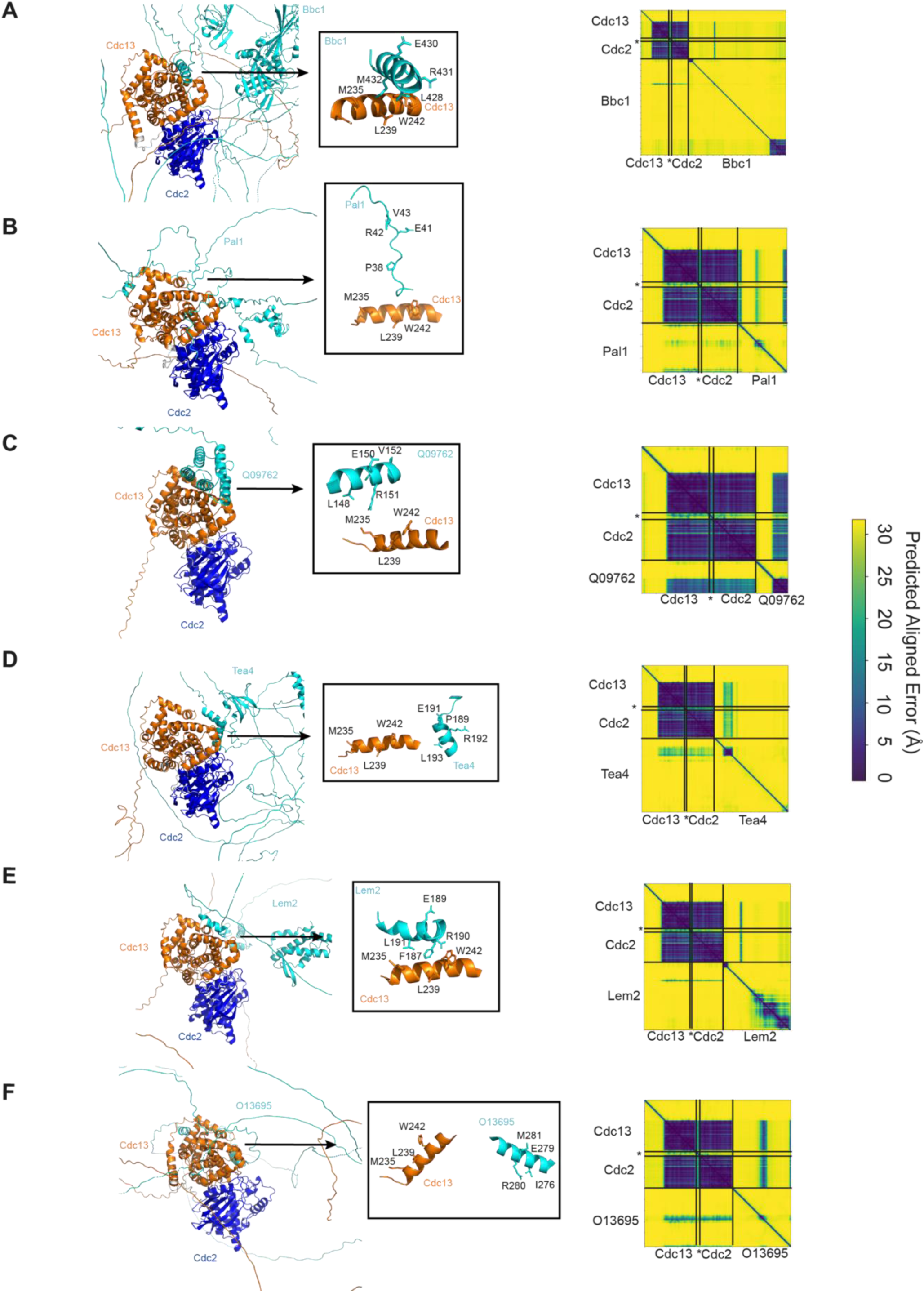
ERL motif-containing proteins based on AlphaPulldown screen. **A-F** Predicted structures of substrates identified to share ERL motifs with Cdc13-L-Cdc2. Box on the right = Cdc13 hydrophobic patch (M235, L239, W242) predicted to interact with the substrates’ respective ERL motif. Right = Respective PAE plot. Black lines indicate boundaries of proteins and internal fusion junctions within the chimeric Cdc13-L-Cdc2 construct. * = Linker of Cdc13-L-Cdc2. **A** Bbc1 **B** Pal1 **C** Q09762 **D** Tea4 **E** Lem2 **F** O13695

**Figure S2:**
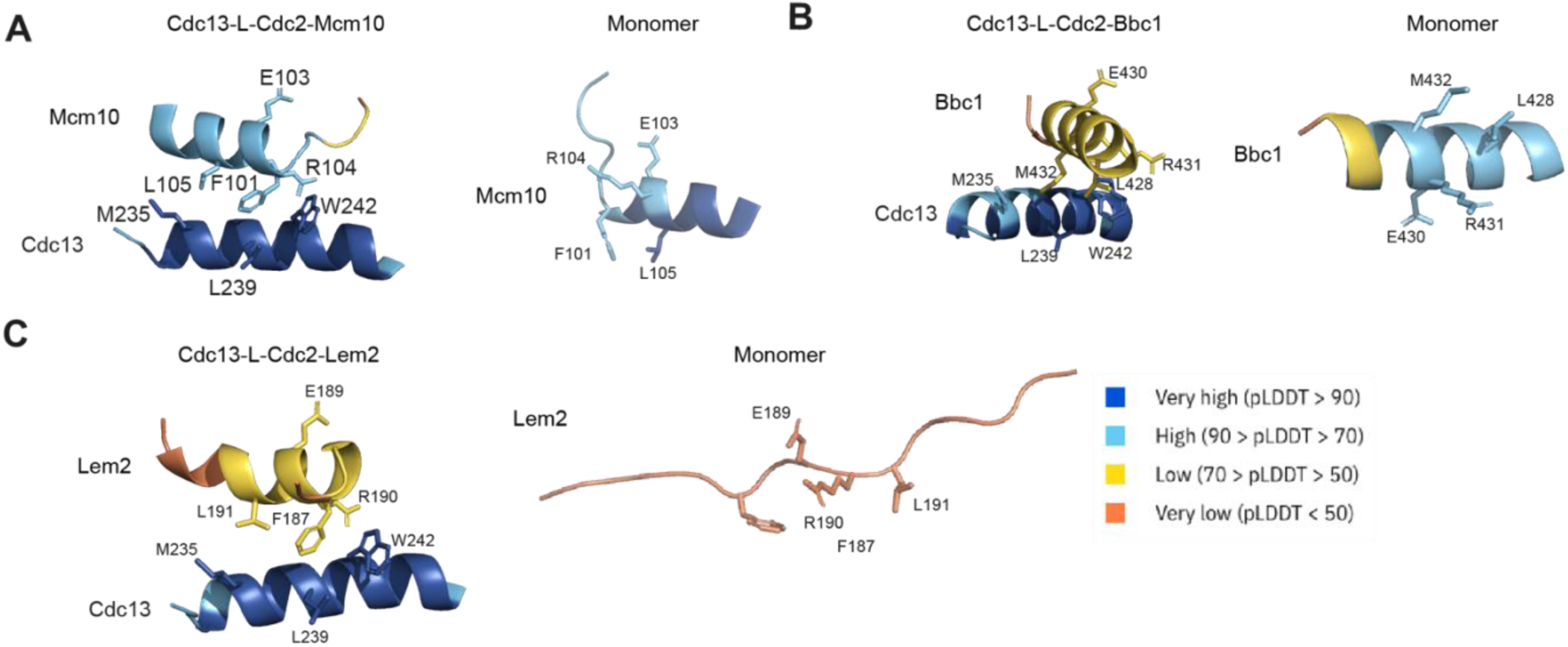
pLDDT of ERL motifs of CDK substrates. **A-C** Left = Cdc13 hydrophobic patch (M235, L239, W242) predicted to interact with the substrates’ respective ERL motif. Right = Monomer structure predictions of CDK substrates with ERL motifs predicted to interact with the Cdc13 hydrophobic patch. Colour indicates pLDDT (Predicted Local Distance Difference Test) = a per-residue measure of AlphaFold’s confidence in local structural accuracy. Monomer structures were obtained from AlphaFold Protein Structure Database. **A** Mcm10 **B** Bbc1 **C** Lem2

**Figure S3:**
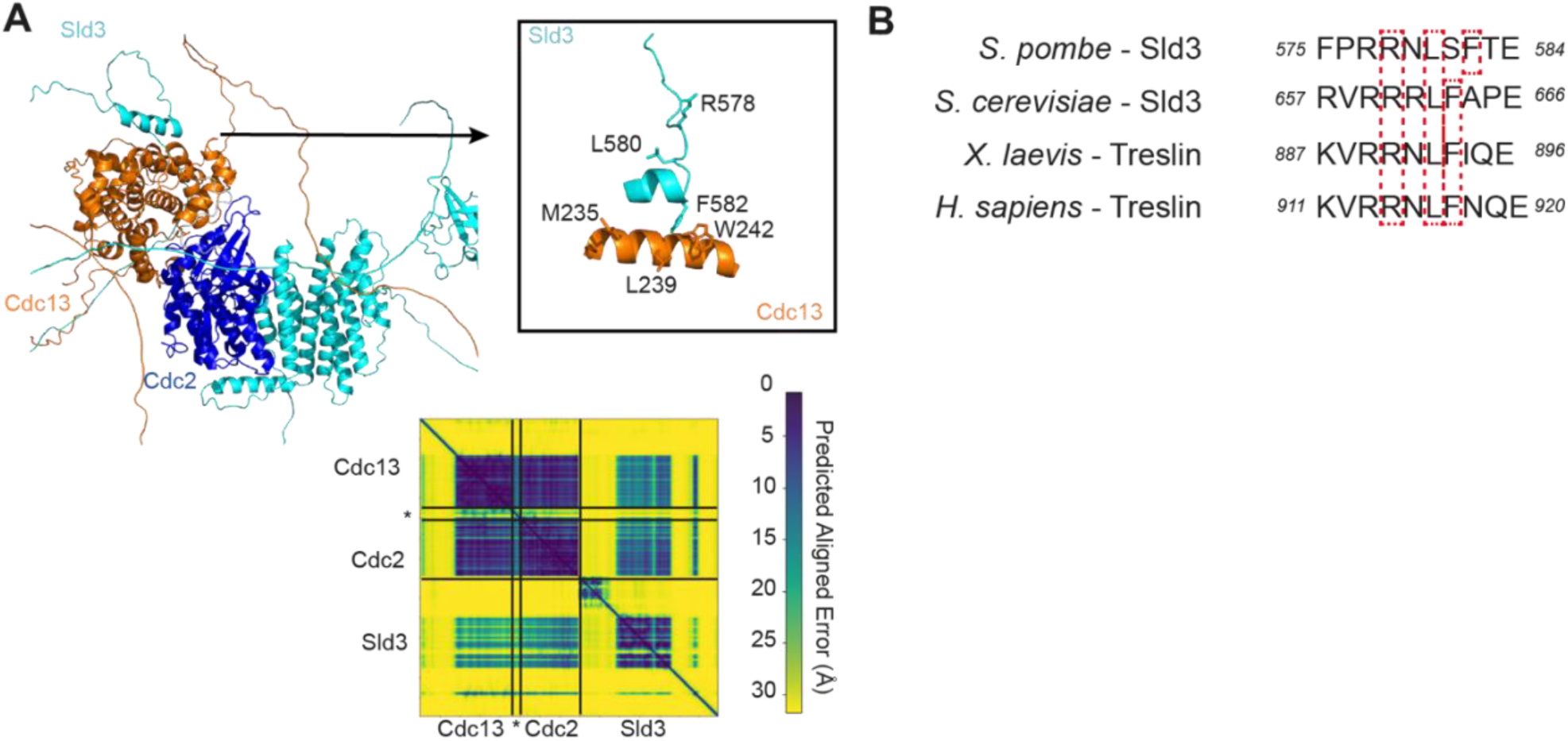
Sld3 conserved RxL motif is predicted to be close to the Cdc13 hydrophobic patch. **A** Top = Predicted structure of Cdc13-L-Cdc2 with Sld3. Box on the right = Cdc13 hydrophobic patch (M235, L239, W242) predicted to interact with RxL motif on Sld3 (R578, L580, F582). Bottom = corresponding PAE plot. Black lines indicate boundaries of proteins and internal fusion junctions within the chimeric Cdc13-L-Cdc2 construct.* = Linker of Cdc13-L-Cdc2. **B** Alignment of Sld3/Treslin RxL(x)F motif and surrounding residues in *S. pombe*, *S. cerevisiae*, *X. laevis* and *H. sapiens*.

**Figure S4:**
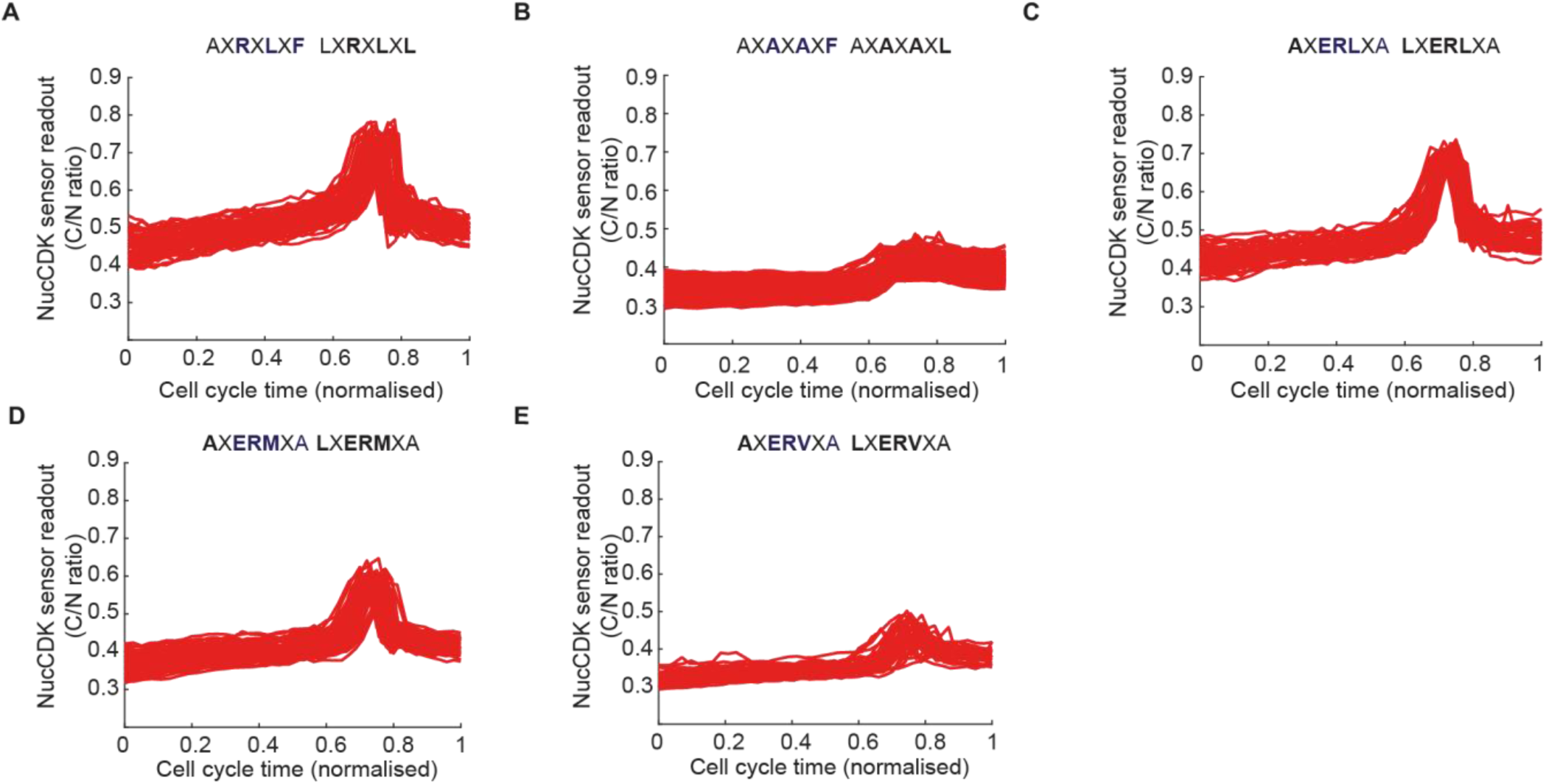
NucCDK traces are consistent between cells. Multiple cell traces of sensors represented in Figure 3 (n > 50 per strain). 1 of 2 biological replicates for each strain is shown. **A** Fig. 3B **B** Fig. 3C **C** Fig. 3D **D** Fig. 3E **E** Fig. 3F

**Figure S5:**
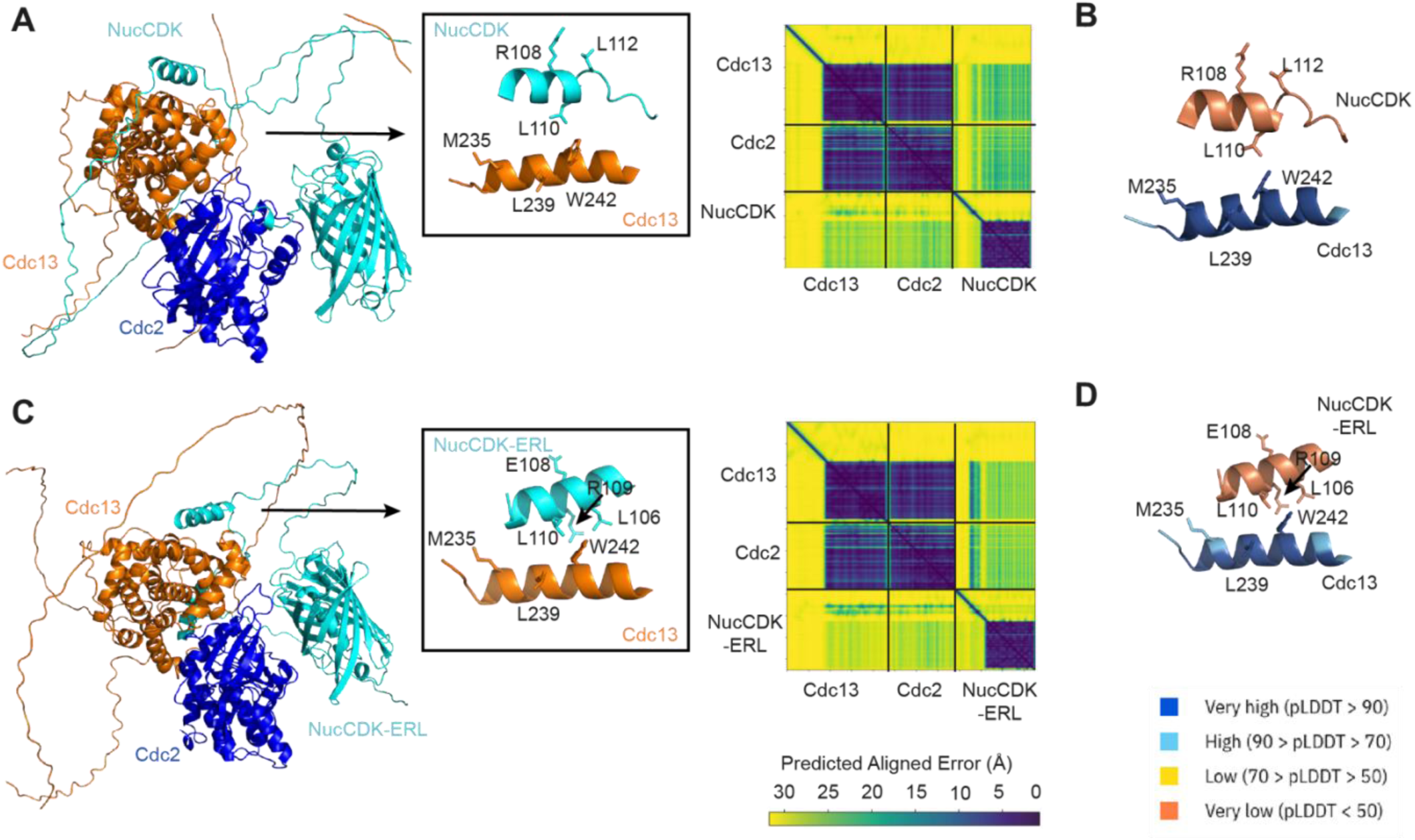
Predicted interaction of NucCDK with Cdc13-Cdc2. AlphaFold-Multimer predicted structures of Cdc13-Cdc2 with **A** NucCDK (ranked_0) **C** NucCDK-ERL motif version (ranked_9). Box on the right = Cdc13 hydrophobic patch (M235, L239, W242) predicted to interact with RxL motif or ERL motif. Right = corresponding PAE plot. Black lines indicate boundaries of proteins. **B & D** pLDDT coloured predicted structures shown in the boxes in **A & C,** respectively.

**Fig. S6:**
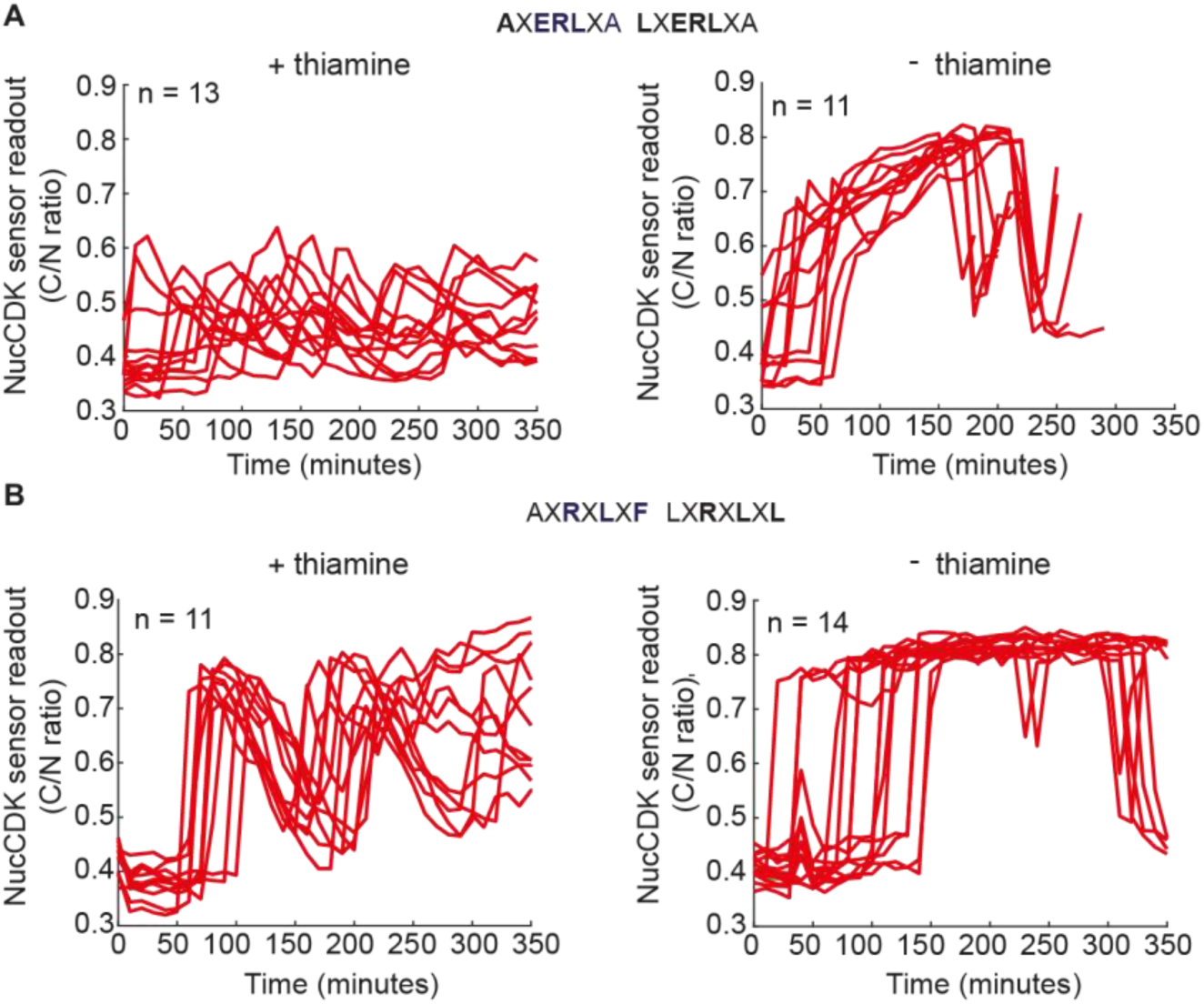
ERL-motif version of NucCDK is phosphorylated less than original NucCDK in the absence of Cdc13. **A** Multiple traces of the experiment in Figure 4B. 1 of 2 biological replicates. **B** Multiple traces of the experiment in Figure 4C. 1 of 2 biological replicates.

